# Three-dimensional human axon tracts derived from cerebral organoids

**DOI:** 10.1101/253369

**Authors:** D. Kacy Cullen, Laura A. Struzyna, Dennis Jgamadze, Wisberty J. Gordián-Vélez, James Lim, Kathryn L. Wofford, Kevin D. Browne, H. Isaac Chen

## Abstract

Reestablishing cerebral connectivity is a critical part of restoring neuronal network integrity and brain function after trauma, stroke, and neurodegenerative diseases. Creating transplantable axon tracts in the laboratory is a novel strategy for overcoming the common barriers limiting axon regeneration *in vivo*, including growth-inhibiting factors and the limited outgrowth capacity of mature neurons in the brain. We describe the generation and phenotype of three-dimensional human axon tracts derived from cerebral organoid tissue. These centimeter-long constructs are encased in an agarose shell that permits physical manipulation and are composed of discrete cellular regions spanned by axon tracts and dendrites, mirroring the separation of grey and white matter in the brain. Features of cerebral cortex also are emulated, as evidenced by the presence of neurons with different cortical layer phenotypes. This engineered neural tissue has the translational potential to reconstruct brain circuits by physically replacing discrete cortical neuron populations as well as long-range axon tracts in the brain.

**eTOC Blurb:** Restoring axonal connectivity after brain damage is crucial for improving neurological and cognitive function. Cullen, et al. have generated anatomically inspired, three-dimensional human axon tracts projecting from cerebral organoids in a transplantable format that may facilitate the reconstruction of large-scale brain circuits.

**Highlights:**

- A neural tissue engineering approach is applied to human cerebral organoids.
- Three-dimensional axon tracts are generated in a transplantable format.
- The growth characteristics of the engineered axons are examined.
- The cellular phenotypes of the organoid tissue and axons are characterized.

## Introduction

Recent advances in understanding the brain from a network perspective (Bassett et al., 2017) suggest that restoring connectivity among different regions of the brain after cerebral injury is crucial for recovering function. However, axon regeneration in the central nervous system is severely restricted after traumatic brain injury, stroke, and other similar conditions because of the presence of environmental axon growth inhibitors (Yiu and He, 2006) and the limited regenerative potential of mature neurons (Fernandes et al., 1999; He and Jin, 2016). Moreover, while intrinsic plasticity mechanisms exist in the brain (Chen et al., 2014), the extent to which the brain can rewire itself is constrained, especially in cases of extensive brain damage. There is thus a critical need to develop strategies for reconstructing brain circuitry.

Prior approaches for restoring axonal pathways include dampening the effects of axon growth inhibitors (GrandPre et al., 2002; Wiessner et al., 2003), augmenting intrinsic neuronal growth programs (Sun et al., 2011), providing pathways that facilitate axon growth (David and Aguayo, 1981; Martinez-Ramos et al., 2012), and adding new neural elements capable of extending processes (Espuny-Camacho et al., 2013; Gaillard et al., 2007). These strategies are all based on the fundamental concept of promoting axon growth *in vivo*. A different approach is the transplantation of preformed axon tracts that are engineered *in vitro*. This alternative relies upon graft integration via the formation of local synaptic connections between transplanted neurons and the host brain but dispenses with the need for long-range axon regeneration. Other benefits include a greater degree of control over and a broader array of available tools for promoting axon growth (Chen et al., 2016). Available technologies for axon engineering include stretch growth (Pfister et al., 2004), patterned substrates (Pan et al., 2015; Smith et al., 1999), and hydrogel micro-columns (Cullen et al., 2012; Struzyna et al., 2015b). Thus far, these methods have utilized primarily dissociated neuronal cultures.

Cerebral organoids are neural tissues derived from the self-organization of pluripotent stem cells (Kadoshima et al., 2013; Lancaster et al., 2013; Pasca et al., 2015; Qian et al., 2016). They develop a significant degree of brain architecture, including rudimentary cortical layers, but large-scale axon bundles are not seen. Inducing the growth of axons from organoids in a controlled manner could lead to repair candidates that replace not only white matter pathways but also grey matter structure (Figure 1A). Here, we combined cerebral organoids with hydrogel microcolumns to generate centimeter-long human axon tracts in a three-dimensional (3D), transplantable format. We report the growth characteristics of neurites in these constructs, as well as the phenotype and structure of the axonal and somatic regions.

**Figure 1.**
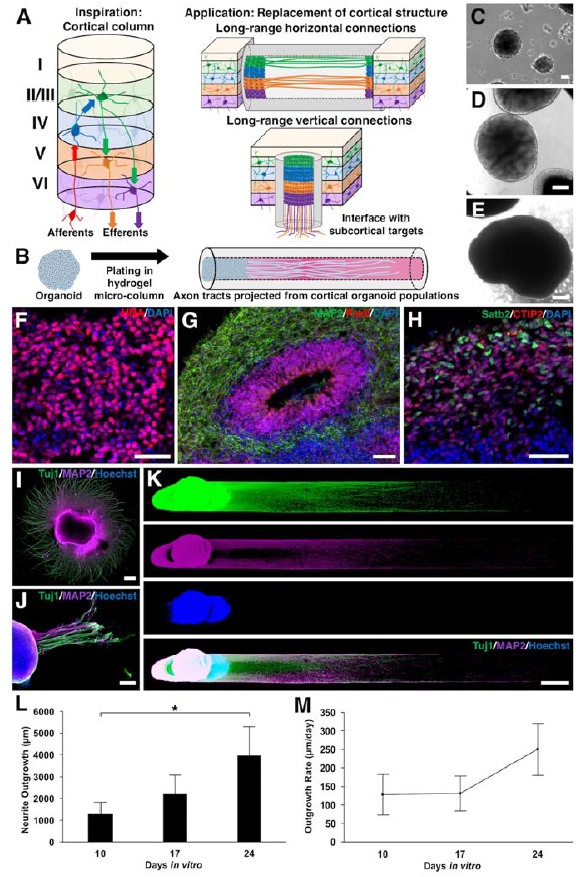
Organoid µTENN generation and unidirectional neurite growth characterization. (A)Cerebral cortex has a stereotyped laminar architecture and vertical circuitry (left). In brief, thalamic inputs enter layer IV, which transmits information to layers II and III. Processed data i s then conveyed from these layers to layers V and VI, which send outputs to subcortical targets. Horizontal projections exist primarily in layers II/III and V. Engineered cortical tissue/axon s could serve as replacements for both long-range horizontal and vertical connections (right). (B) The H9 human embryonic stem cell line was used to generate cerebral organoids using a modified version of existing protocols (Pasca et al., 2015). Between dd100-150, cut pieces of oragnoids were inserted into one or both ends of the µTENN micro-column. The micro-column lumen provided the necessary directionality for the formation of aligned neurite tracts. Phase contrast micrographs show H9 embyroid bodies (C), and dd15 (D) and dd45 cerebral organoids (E). (F) Immunofluorescence staining confirmed the human origin of these organoids (human nuclear antigen: red). (G) Organoids (dd61) recapitulated neurodevelopmental structures, including ventricle-like zones surrounded by neural progenitors (Pax6: red) and differentiated neurons (MAP2: green). (H) Separation of upper‐ (Satb2: green) and lower-layer (CTIP2: red) cortical neurons was also observed (dd61). (I) Organoids plated on a planar surface extend neurites in an isotropic manner (MAP2: magenta, Tuj1: green, Hoechst: blue). (J-K) In contrast, organoids inserted into a µTENN micro-column displayed directional growth of neurites within the micro-column lumen. J shows a construct that had extruded from the agarose outer shell (J: 25 DIV, K: 24 DIV; MAP2: magenta, Tuj1: green, Hoechst: blue). Neurite outgrowth distances (L) and growth rates (M) were quantified in unidirectional organoid µTENNs cultured over 10, 17 and 24 days *in vitro*. Data are presented as mean ± standard deviation with * denoting statistical significance (*p* < 0.05). Scale bars: 500 µm (C, E, I, K), 200 µm (D), 40 µm (F), 100 µm (G), 50 µm (H), and 150 µm (J).

## Results

### Generation of organoid µTENNs

Micro-tissue engineered neuronal networks (µTENNs) are hydrogel columns that are hundreds of microns in diameter and patterned to separate neuronal somata from aligned neurites (Cullen et al., 2012; Struzyna et al., 2015a). This configuration recreates, in part, the segregation of neuronal populations and long-range axon tracts found in the brain. These constructs consist of a relatively firm hydrogel shell that can withstand physical manipulation and an extracellular matrix core conducive to cellular and neurite growth (Figure 1B). In the present study, we engineered agarose tubes containing a cross-linked type I collagen core that facilitated the directed growth of neurites.

Cortical organoids were generated from the H9 human embryonic stem (ES) cell line using a modified version of a previously published protocol (Pasca et al., 2015). Embryoid bodies derived from enzymatic elevation of whole ES cell colonies (Figure 1C) developed into organoids with multiple internal substructures and peripheral translucency, a marker of developing neuroepithelium (Figures 1D-E). These organoids expressed human nuclear antigen (Figure 1F) and formed ventricle-like structures surrounded by an adjacent layer of neural progenitors (Pax6+) and an outer layer of differentiated neurons (MAP2) (Fig. 1G). By differentiation day (dd) 61, both upper‐ (Satb2+) and lower-layer (CTIP2+) cortical neurons were identified with the development of rudimentary laminar architecture (Figure 1H). While organoids cultured on a planar surface extended neurites in an isotropic fashion (Figure 1I), organoid tissue inserted into one end of a hydrogel micro-column projected a combination of axons (Tuj1+/MAP2-) and dendrites (Tuj1+/MAP2+) exclusively along the length of the collagen matrix core (Fig. 1J).

Taking advantage of this directional growth of organoid-derived neurites, we generated unidirectional organoid µTENNs (Figure 1K) and determined their neurite growth characteristics over 24 days *in vitro* (DIV). Of note, Hoechst labeling revealed little evidence of neuronal migration into the collagen core, resulting in a relatively pure segment of neurites. Quantification of neurite outgrowth yielded a length of 3,995±1,356 µm (mean ± standard deviation; Figure 1L). The neurite growth rate accelerated to reach 250±69 µm/day by 24 days DIV (Figure 1M).

### Neurite segment characterization of bidirectional organoid µTENNs

We next assessed the structural details and growth patterns of neurites in bidirectional µTENNs, in which organoid tissue was inserted into both ends of the hydrogel micro-column. Green-fluorescent protein (GFP)-positive organoids (Paluru et al., 2014) were used to assist with visualization of neurites. Initially, 0.5 cm constructs were generated (Figure S1A). At this construct length, neurites crossed the entire length of the collagen core by 24 DIV (Figure S1C). To quantify neurite densities, the normalized mean fluorescence intensity of 5 different regions of interest (ROI) in the neurite segment was measured (Figure S1A, S1D). This analysis showed that both GFP and Tuj1 intensity ratios were similar in all ROI’s that were not immediately adjacent to the organoid cell mass, suggestive of a neurite network that had reached a steady-state density by 24 DIV.

At a longer construct length of 1 cm, neurites from each side grew a considerable distance by 24 DIV but had not fully crossed at the center of the neurite segment (Figure 2A). Continued neurite growth resulted in what appeared to be the formation of a unified neurite network by DIV 60 (Figure 2B). On a qualitative basis, neurite density at the center of the neurite segment increased over time. Individual neurites could be resolved near the center of the µTENN at DIV24 (Figure 2C), but neurite density increased considerably in the same region by DIV60 (Figure 2D). To quantify this observation, we again measured normalized mean fluorescence intensities across 5 ROI’s in the neurite segment. At 24 DIV, GFP and Tuj1 intensity ratios progressively decreased from the peripheral to the central parts of the neurite segment (Figure 2E). At 60 DIV, Tuj1 intensity ratios were similar across all ROI’s, and GFP intensity ratios were similar between the inner and central ROI’s. These results suggested active neurite growth and remodeling at 24 DIV and a steady-state neurite density at 60 DIV similar to the 24 DIV results for 0.5 cm constructs. Of note, the neurites found throughout 0.5 and 1 cm constructs at both time points consisted of a combination of axons (NF200+/MAP2-) and dendrites (NF200+/MAP2+) (Fig 2F).

**Figure 2.**
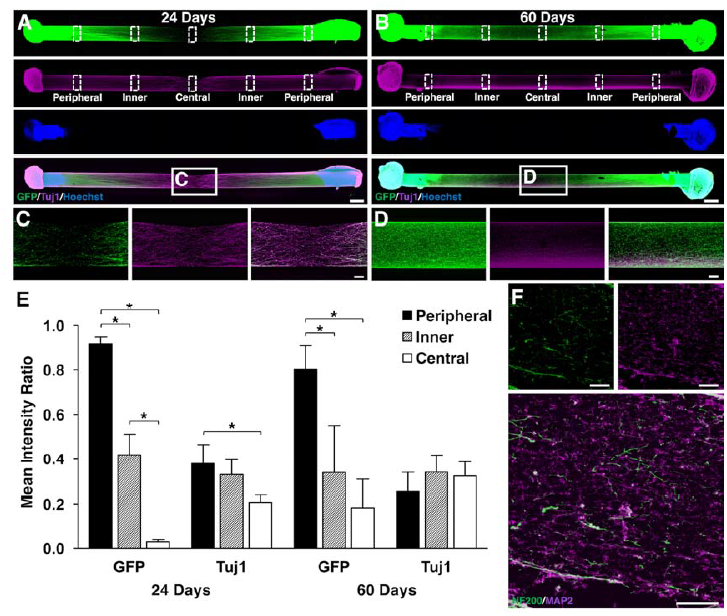
Phenotypic and growth characterization of neurites in 1 cm bidirectional μTENNs. Confocal reconstructions of 1 cm GFP-expressing constructs cultured for 24 (A) or 60DIV (B) and stained for Tuj1 (magenta) and Hoechst (blue). Higher magnification images of th neurite tracts qualitatively show that neurites populate the center of the micro-column with a lesser density at 24 DIV (C) compared to 60 DIV (D). (E) Quantification of the mean GFP and Tuj1 intensities in the different regions of interest in A and B confirm that neurites at the center of the µTENN reach a steady-state density at the later time point. Data are presented as mean ± standard deviation with * denoting statistical significance (*p* < 0.05). (**F**) Confocal image of th neurite tract in a sectioned organoid µTENN shows expression of both neurofilament 200(NF200; green) and microtubule-associated protein 2 (MAP2; magenta), indicating the presence of both axons and dendrites. Scale bars: 500 µm (A-B), 100 µm (C-D, F).

### Somatic segment characterization of bidirectional organoid µTENNs

At both construct lengths, Hoechst staining again confirmed the absence of neuronal migration into the collagen core (Figures S1A and 2A-B). Notably, the area of Hoechst staining increased substantially between the two time points, which indicated continue growth of the organoid tissue (Figure 3A). Higher magnification examination of the organoid tissue at both time points revealed the formation of cellular aggregates with radial processes, particularly in Tuj1 stains (Figures S1B and 3B). This pattern of cell clustering is not typically seen in conventional organoids but can be encountered in planar and 3D hydrogel cultures of dissociated neurons.

Additional immunohistochemical analysis was performed to examine the cellular composition of the organoid tissue. Astrocytes (GFAP+) were found throughout the somatic segment at both time points (Fig. 3C-D). Pax6+ cells were also scattered throughout the somatic segment, which indicated the continued presence of some progenitor cells at time points greater than differentiation day (dd) 120 (Figure 3E) and explained the growth of the cell mass seen in Figure 3A. The majority of the cells within this region stained positive for Tuj1 (Figure 3C), confirming their neuronal identity.

**Figure 3.**
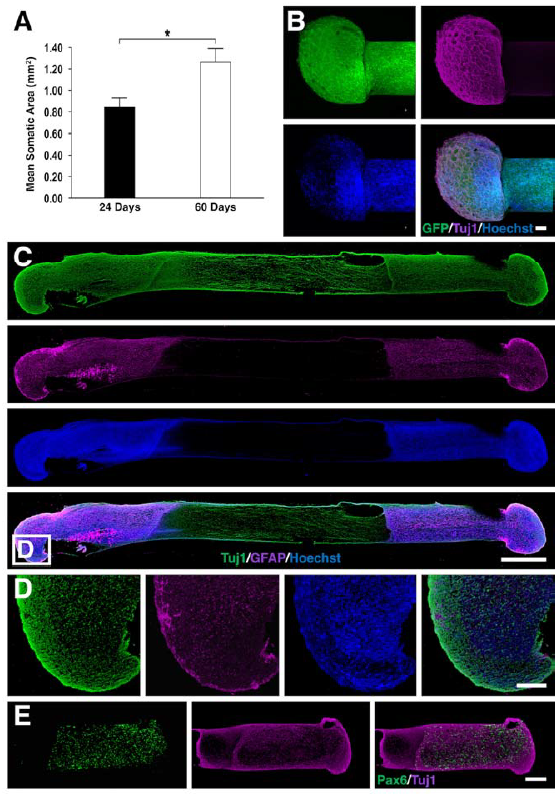
Characterization of the somatic segment of the organoid μTENNs. (A) The mean cell mass area was quantified from Hoechst staining of 1-cm µTENNs at 24 and 60 DIV. There was a significant increase in somatic area as a function of time. Data are presented as mean ± standard deviation with * denoting statistical significance (*p* < 0.05). (B) High-magnification image of the organoid cell mass from a 1-cm GFP-expressing µTENN at 24 DIV stained with Tuj1 (magenta) and Hoechst (blue) showing neuronal clusters connected by neurite tracts in a radial configuration. (C) Full-length confocal reconstruction of a 12-µm thick section of a 0.5 cm µTENN at 24 DIV stained for Tuj1 (green), GFAP (magenta) and Hoechst (blue). Astrocytes are scattered throughout the organoid aggregate. (D) Magnified images of the aggregate region referenced in the box inset in C. (E) Pax6+ neural progenitors are found throughout the organoid cell mass (1 cm construct at 24 DIV). Scale bars: 500 µm (A), 100 µm (B-D), and 150 µm (E).

Neurons positive for Satb2 and CTIP2, which are markers for primarily layer II/III (callosal) and layer V cortical neurons, respectively (Molyneaux et al., 2007), were identified within the organoid tissue. No structural organization could be identified in some cases (Figure 4A), but some degree of laminar structure was preserved in others (Figure 4B). Overall, constructs lacking structure were more commonly observed (Figure 4C). However, the frequency of constructs with segregation of Satb2+ and CTIP2+ cells increased from 24 DIV to 60 DIV (Figure 4C-D). Co-localization of Satb2 and CTIP2 labeling was observed at both time points (Fig. 4E), a finding that has been associated with a distinct subpopulation of cortical projection neurons (Harb et al., 2016). The percentage of CTIP2+ cells also expressing Satb2 decreased from 24 DIV to 60 DIV, which mirrors results from murine development (Harb et al., 2016; Leone et al., 2015).

**Figure 4.**
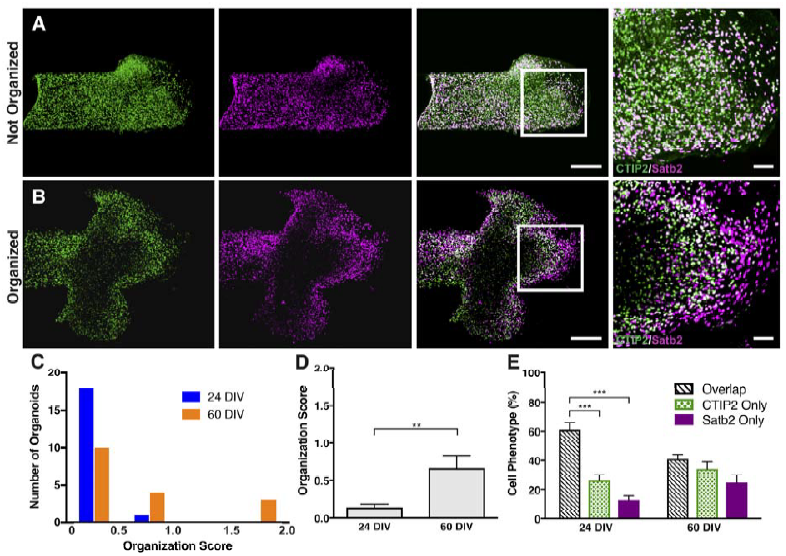
Satb2 and CTIP2 organization and co-localization. (A-B) Confocal images of two different µTENN constructs demonstrate the presence of upper‐ (Satb2, magenta) and lower - layer (CTIP2, green) cortical neurons in the organoid cell mass. Intermixing of these two populations is observed in A while some degree of laminar segregation is seen in B. Scale bars: 200 µm (A-B main), 50 µm (A-B inset). (C) Each side of a cohort of constructs (24 DIV: n=11 constructs/19 available sides, 60 DIV: n=10 constructs/17 available sides) was scored for its degree of structural organization based on the spatial distribution of Sabt2+ and CTIP2+ cells (0=no organization, 1=geographic segregation but no laminar structure, 2=laminar structure). Averaged scores from three blinded authors (DKC, WG, HIC) are depicted as histograms with 4 bins (≤0.5, >0.5 and ≤1, >1 and ≤1.5, and >1.5 and ≤2) for each time point. (D) The mean of the averaged organization scores for each time point is illustrated. (E) In the same group o fconstructs that was analyzed for structure, the frequency of Satb2+, CTIP2+, and Satb2+/CTIP2+ cells was quantified at each time point. Data in D and E are presented as mean  SEM (**p<0.01, ***p<0.001).

## Discussion

We have produced the first example of tissue-engineered, long-projecting cortical axon tracts derived from human stem cell sources. These constructs mirror the architecture of grey and white matter in the brain and are designed to tolerate the physical manipulations required for transplantation into the brain. By combining principles from the disciplines of stem cell biology and neural tissue engineering (Yin et al., 2016), this work opens the possibility of rewiring the brain with patient-specific, lab-grown laminar cortical tissue spanned by axon tracts.

Developing methods for repairing axonal injury in the brain is particularly important because of the vital role axons serve in maintaining cerebral network function (Follett et al., 2009; Riley et al., 2011). The approach of generating axon tracts in the lab has certain advantages over promoting axon growth *in vivo*, including greater control over cell types, gene expression, and growth conditions (Chen et al., 2016). At the time of transplantation, there is also greater control over which regions of the brain are reconnected. The concept of µTENNs is one example of this approach that permits precise control over the architecture and physical properties of a 3D construct (Cullen et al., 2012; Harris et al., 2016). Previously, it has been demonstrated that µTENNs built from aggregates of dissociated rat embryonic cortical neurons survive, maintain their structure, and project neurites into adjacent brain tissue up to 1 month after stereotactic implantation (Struzyna et al., 2015b). The current work improves upon the translational potential and increases the sophistication of µTENN technology by utilizing cerebral organoids as the source of neurons and neurites. These structured neural tissues can be derived from patient-specific induced pluripotent stem (iPS) cell lines, and they exhibit the architecture and possibly the micro-circuitry of laminar cortex that is lacking in re-aggregated neuron clusters (Birey et al., 2017; Kadoshima et al., 2013; Lancaster et al., 2013; Pasca et al., 2015; Qian et al., 2016).

Several areas of improvement would benefit future iterations of organoid µTENNs. Although we observed segregation of cortical layer-specific markers in some constructs, most showed intermixing of these markers. This lack of structure could result from variability in organoid growth (Quadrato et al., 2017) or organoid manipulation during µTENN generation. Regarding the latter point, we found that the frequency of organized constructs increased over time. One potential explanation for this observation is that the physical disruption caused by organoid manipulation was mitigated in some cases by continued organoid growth, which restored structure through intrinsic programs of self-organization. The lack of or incomplete restoration of structure could be the product of insufficient time or a degree of structural disruption that was too great to overcome. Prior studies have documented that up to 40% of CTIP2+ cells also express Satb2 (Alcamo et al., 2008). We found a higher percentage of CTIP2+ cells co-localizing with Satb2, which could result from differences in species (human versus rodent), time points examined, and choice of antibodies. Improved control of organoid architecture and phenotype will be important for facilitating the creation of custom cortical constructs, such as vertical versus horizontal circuits (Figure 1A).

Our current protocol for creating human cortical axon tracts 1 cm in length from organoid tissue required 2-3 months for organoid preparation and 1-2 months for axon growth. Generating longer axon lengths in shorter periods of time would be important for human translation. Younger organoids or methods for accelerating organoid maturation (Li et al., 2017) could be used to shorten the time required for organoid preparation. Faster axon growth rates could be attained using growth factor regimens or by adapting other bioengineering techniques, such as axon growth via mechanical stretch (Pfister et al., 2004).

Applications of organoid µTENNs extend beyond reconstructing brain circuitry to serving as physiologically relevant models for studying internodal network function in human neurodevelopment and disease. This three-dimensional system more faithfully replicates the modular structure of the brain than dissociated planar cultures (Shein-Idelson et al., 2011; Shein-Idelson et al., 2016), and they possess axon tracts that are not found in standard cerebral organoids. Moreover, organoid µTENNs may be more relevant to human disease than animal models (Lancaster and Knoblich, 2014). Using this platform to study network function will require further examination of the intrinsic neuronal activity of cerebral organoids (Birey et al., 2017; Quadrato et al., 2017) and detailed investigation of the capacity of organoid µTENNs for transmitting data across its axons (Chen et al., 2017). An understanding of how activity propagates across organoid µTENNs also would help establish the efficacy and limitations of using these constructs to restore axonal connectivity *in vivo*.

## Experimental Procedures

### Embryonic stem cell maintenance

The H9 human embryonic stem cell line expressing enhanced green fluorescent protein (Paluru et al., 2014) was maintained on mouse embryonic fibroblast feeder cells. Stem cells were cultured in human embryonic stem cell (hES) media, and colonies were passaged every 5-6 days. Passage numbers were between 110-135.

### Generation of cerebral organoids

Cerebral organoids were generated using a protocol modified from Pasca, et al., 2015. In brief, stem cell colonies were detached with dispase. Embryoid bodies were then placed in ultra-low attachment 12-well plates in Induction Media (hES media supplemented with 100 nM LDN193189, 10 µM SB431542, 2 µM XAV939, and 2.5 µM Y-27632). From differentiation day (dd) 1-6, the developing embryoid bodies were maintained in Induction Media without Y-27632. From dd6-25, the developing organoids were maintained in Neuronal Media (Neurobasal media supplemented with 1:50 B27 without vitamin A, 1X Glutamax, 100 U/mL penicillin/100 µg/mL streptomycin, 20 ng/mL bFGF, and 20 ng/mL EGF). From dd25-43, media consisted of Neuronal Media supplemented with 20 ng/mL NT3 and 20 ng/ML BDNF. After dd43, Neuronal Media without EGF and bFGF was used. Organoids used for µTENN construction were between dd100-150.

### Fabrication of micro-column constructs

The outer hydrogel shell of the micro-columns, which consisted of 1% agarose (outer diameter of 973 µm), was generated by drawing the agarose solution into a capillary tube via capillary action. An acupuncture needle (500 µm diameter) was inserted into the center of the agarose-filled capillary tube before agarose polymerization to produce space for an inner core. The inner core consisted of a rat-tail type I collagen solution. Organoids were cut into small pieces using fine-tip forceps and inserted into one or both ends of the micro-columns. In some cases, the organoids were subsequently trimmed with forceps to fit fully within the micro-column. Organoid µTENN cultures were cultured in Neuronal Media without EGF or bFGF.

### Immunohistochemical analysis of organoid µTENNs

Whole constructs or paraffin-embedded sections were fixed in 4% paraformaldehyde, and the latter underwent dewaxing and antigen retrieval procedures. Following permeabilization and blocking steps, samples were incubated with primary antibodies (see Table S1) and appropriate secondary antibodies.

### Data quantification and statistical analyses

The length of neurite outgrowth in the unidirectional constructs was determined by measuring the longest observable neurite in each construct from the organoid cell mass. The growth rate and corresponding standard deviation at each timepoint were estimated by the backward difference method using the mean axonal outgrowth and its standard deviation, respectively. To compare the neurite density along the micro-columns, five regions of interest (ROI) of equal size were selected in the GFP and Tuj1 channels along the region spanned by axon tracts. The ROIs were corrected for background and normalized to the maximum value of the organoid cell mass region of each sample. ImageJ was used to measure the area of the organoid cell mass in the Hoechst channel. Counting of Satb2+, CTIP2, and Satb2+/CTIP2+ cells were performed using a custom automated algorithm, which was validated by manual counting. Organoid tissue architecture was qualitatively categorized using a 0-2 scale (0=no organization, 1=geographic segregation of Satb2+ and CTIP2+ cells but no laminar structure, 2=laminar structure).

## Acknowledgments

Financial support was provided by the National Institutes of Health [U01-NS094340 (Cullen) and T32-NS091006-02S1 (Struzyna)], Department of Veterans Affairs [Career Development Award #IK2-RX002013 (Chen) and Merit Review #B1097-I (Cullen)], Michael J. Fox Foundation [Therapeutic Pipeline Program #9998 (Cullen)], and the Neurosurgery Research and Education Fund [Bagan Family Young Clinician Investigator Award (Chen)]. The authors thank Saarang Karandikar for assistance in data analysis.

Reported data is available upon request. DKC and HIC conceived this project and provided general supervision. DJ and JL maintained stem cell cultures and generated cerebral organoids. LAS generated organoid μTENNs. LAS, WJG, KLW and KDB performed immunohistochemical analysis and/or associated data analysis. All authors contributed to drafting and editing the manuscript.

